# Mesophyll conductance in two cultivars of wheat (*Triticum aestivum*) grown in glacial to super-elevated [CO_2_]

**DOI:** 10.1101/2020.04.29.069492

**Authors:** Eisrat Jahan, Peter C. Thomson, David T. Tissue

**Author notes:** Author for correspondence: Dr Eisrat Jahan, The University of Sydney.

## Abstract

Mesophyll conductance (*g*_m_) is an important factor limiting photosynthesis. However, *g*_m_ response to long-term growth in variable [CO_2_] is not well understood, particularly in crop plants. Here, we grew two cultivars of wheat (Halberd and Cranbrook), known to differ in *g*_m_ under current environmental conditions, in four [CO_2_] treatments: glacial (180 μmol mol^−1^), pre-industrial (280 μmol mol^−1^), current ambient (450 μmol mol^−1^) and super-elevated (1000 μmol mol^−1^) in well-watered and moderate water limitation conditions, to develop an evolutionary and future climate perspective on *g*_m_ control of photosynthesis and water use efficiency (WUE). In the two wheat genotypes, *g*_m_ increased with rising [CO_2_] from glacial to ambient [CO_2_], but declined at super-elevated [CO_2_]; however, the specific mechanism of *g*_m_ response to [CO_2_] remains unclear. Although *g*_m_ and *g*_m_/*g*_sc_ (mesophyll conductance/stomatal conductance) were strongly associated with the variability of *A* and WUE, we found that plants with higher *g*_m_ may increase *A* without increasing *g*_sc_, which increased WUE. These results may be useful to inform plant breeding programs and cultivar selection for Australian wheat under future environmental conditions.

**Highlight:** Mesophyll conductance increased with increasing [CO_2_] from glacial to ambient CO_2_ levels, then declined at super-elevated CO_2_ for both well-watered and water-limited treatments. These responses of mesophyll conductance with varying [CO_2_] have a physiological basis.

## Introduction

Climate change is a major environmental challenge with significant threats to water resources, crop development and food security. CO_2_ concentration has increased by 47% (278 ppm to 408 ppm) from 1750 to 2018 (IPCC, 2019) and is predicted to reach 550 μmol mol^−1^ in the middle of this century and 700 μmol mol^−1^ by the end of this century (Prentice *et al*., 2001). This rising atmospheric CO_2_ concentration is altering global temperature and precipitation patterns, which challenges agricultural productivity (Ainsworth *et al*., 2008). From the perspective of food security, a major uncertainty is the extent to which crops will serve as a sink for extra atmospheric CO_2_, and if this will affect food nutritive quality (Prentice *et al*., 2001). Therefore, a major challenge for plant physiologists, agronomists and breeders is to select appropriate physiological traits that can maximize future crop production and quality in a challenging climate.

Increasing CO_2_ concentration generates partial closure of stomata, thereby decreasing leaf conductance to CO_2_ and H2O vapour (Morison, 1985, 1987; Allen, 1990) and reducing leaf transpiration (Kimball and Idso, 1983). This reduced stomatal conductance can increase leaf temperature and potentially negate decreases in evapotranspiration (ET) (Kimball *et al*., 1992, 1994, 1995). Elevated CO_2_ can reduce plant water consumption, leading to reduced soil water depletion (Hungate *et al*., 1997; Leuzinger and Korner, 2007; Robredo *et al*., 2007) but it also often increases the leaf area index counteracting leaf-level reductions in transpiration rate (Franks *et al*., 2013). In addition, plant-scale water use may not decline in elevated CO_2_. Moreover, in the future, prolonged summer droughts and higher frequency of extreme precipitation events will become more common (IPCC, 2007). Plant water status affects the magnitude of plant carbon uptake in elevated CO_2_ (Volk *et al*., 2000; Knapp *et al*., 1999; Morgan *et al*., 2001). So in the future, elevated CO_2_ combined with water scarcity may drastically affect crop production. Furthermore, the annual global wheat demand will increase to about 900 Mt by 2050 with limited expansion of sown area due to increased population and urbanisation (Alexandratos and Bruinsma, 2012).

During photosynthesis, CO_2_ diffuses from the atmosphere (C_a_) to the sites of carboxylation (C_c_) inside the chloroplasts (Farquhar *et al*., 1980). This CO_2_ uptake is strongly influenced by environmental factors that are predicted to change in the future, namely, atmospheric CO_2_ concentration, temperature and water availability (Sage and Kubien, 2007; Lawlor and Tezara, 2009; Alberth *et al*., 2011). Mesophyll conductance is the conductance of CO_2_ from the leaf intercellular air space (C_i_) to the sites of carboxylation (C_c_), including gas and liquid phase transfer. Mesophyll conductance (*g*_m_) can strongly influence the rate of photosynthesis (Warren *et al*., 2003; Flexas *et al*., 2008; Barbour *et al*., 2010; Giuliani *et al*., 2013; Jahan *et al*., 2014; Xiong *et al*., 2016; Ellsworth *et al*., 2018; Tosens and Laanisto, 2018; Han *et al*., 2018; Knauer *et al*., 2019) with up to 25-30% reduction in photosynthetic rate (Epron *et al*., 1995), and this limitation may be as large as that imposed by stomatal conductance (*g*_sc_) (Warren and Adams, 2006; Flexas *et al*., 2007a). Mesophyll conductance is a dynamic leaf trait that depends on genotype, environmental factors and anatomical differences; *g*_m_ may be affected by drought stress, CO_2_ concentration and salinity during the growing period (Flexas *et al*., 2008).

Leakey *et al*. (2009) reviewed the effect of long-term elevated CO_2_ treatment on growth, and concluded that photosynthetic carbon gain is enhanced in C3 plants by elevated CO_2_ despite partial acclimation of photosynthetic capacity. Singsaas *et al*. (2003) reported that an increase in CO_2_ concentration (200 μmol CO_2_ mol^−1^ above atmospheric level) decreased *g*_m_ for some species [cucumber (*Cucumis sativus* L.), spinach (*Spinacia oleracea* L.) and linden bean (*Phaseolus vulgaris* L. var. Linden)] but not for all. Conversely, Bernacchi *et al*. (2005) showed that mesophyll conductance of soybean (*Glycine max*) was not altered by elevated-growth CO_2_ levels. However, studies into the effects of elevated growth CO_2_ on *g*_m_ are scarce (Singsaas *et al*., 2003). To the best of our knowledge, there is no published report on the variation in mesophyll conductance of wheat with different growth CO_2_ concentrations.

In C_3_ leaves, water stress predominantly affects CO_2_ diffusion in leaves through a decrease of stomatal and mesophyll conductance without decreasing the biochemical capacity to assimilate CO_2_ (Flexas *et al*., 2004). Tosens *et al*. (2012) worked on saplings of *Populus tremula* L. with drought and found that water stress reduced chloroplast surface area to leaf area ratio (*S*_c_/*S*) and increased cell wall thickness and these are partly responsible for lower *g*_m_. In contrast, *g*_m_ was unaffected by drought in bell pepper [*Capsicum annuum* L. var. *anuum* (Grossum Group) ‘Quadrato d’Asti’] (Delfine *et al*., 2001) and sugar beet (*Beta vulgaris* L.) plants (Monti *et al*., 2006). Mesophyll conductance was strongly reduced for grapevines on the first day of water stress, but it was restored to the pre-stress values by the fourth day of water stress (Pou *et al*., 2012). If we consider water stress and elevated CO_2_ together, then existing studies suggest that plant responses to elevated [CO_2_] and drought stress are highly variable (Atwell *et al*., 2007; Domec *et al*., 2010; Wertin *et al*., 2010, 2012; Ayub *et al*., 2011; Duursma *et al*., 2011; Warren *et al*., 2011; Zeppel *et al*., 2012; Franks *et al*., 2013; Lewis *et al*., 2013; Perry *et al*., 2013; Duan *et al*., 2014). In this study, the following hypotheses were tested: A) the historical rise in CO_2_ from glacial to predicted future environment will affect *g*_m_, and the responses of *g*_m_ with varying [CO_2_] have a physiological basis; B) *g*_m_ is one of the dominant factors regulating photosynthesis; and C) *g*_m_ plays an important role in improving photosynthesis and water use efficiency (WUE) simultaneously in a future-stress climate scenario.

## Materials and Methods

### Plant material and growth conditions

Two cultivars (Cranbrook, Halberd) of wheat (*Triticum aestivum* L.) were grown in four controlled-environment growth rooms at the University of Sydney, Centre for Carbon Water and Food (CCWF), Camden (NSW, Australia). The wheat cultivars ‘Halberd’ and ‘Cranbrook’ are parental lines that were studied in depth in a quantitative trait loci (QTL) mapping population studied for Δ^13^C of leaf tissue by Rebetzke *et al*. (2008). These two cultivars also have a wide range of *g*_m_ values under non-stress environmental conditions (Jahan *et al*., 2014). These four growth rooms differed only in their (ambient growth) CO_2_ mole fractions. Growth CO_2_ levels were set to 180, 280, 450 and 1000 μmol mol^−1^ to represent four [CO_2_] from glacial, pre-industrial, ambient and super-elevated periods. Measured [CO_2_] inside growth rooms were 206, 344, 489 and 1085 μmol mol^−1^, using a Picarro ^13^CO_2_ laser analyser (G1101-I, Picarro, CA, USA) to draw air from each room in sequence to the laser for 10 minutes each. Average [CO_2_] and δ^13^*C* of room air (δ^3^ *C_CO2_*) were calculated over the last 5 minutes of sampling for the whole growing period and stopped 7 days prior to measurement. The running average over the 7 days prior to gas exchange measurement was used in *g*_m_ calculations described below, and for Δ^13^C calculation of leaf tissue samples. The laser was calibrated as described by Thurgood *et al*. (2014). Apart from [CO_2_], other environmental characteristics inside all the cabinets were the same: temperature was 25 °C during the 14-h light period (PPFD was 400 μmol m^−2^s^−1^ at the top of the plants), and 17 °C during the 10-h dark period, while relative humidity was 75% at all times.

Seeds were planted in 2-L pots filled with commercial potting mix with slow-release fertilizer (Osmocote Exact, Scotts, Sydney, NSW, Australia). Seedlings were thinned to one per pot after emergence. Sixteen days after emergence water treatments were started by withholding water to half of the pots as a drought treatment. At the temporary wilting point (7 days after the start of the drought treatment), weights were recorded for all drought pots, and this was taken as a target water content. A visual assessment of leaf wilting was used as an indicator of water status for water-stressed plants where temporary wilting point was defined as the first day on which leaves of droughted plants wilted. The target water content was maintained gravimetrically thereafter to introduce a mild water stress to half of total plants. The irrigated pots were well watered throughout the experiment.

### Measurement of g_m_

Two or three fully expanded youngest leaves per plant were selected to measure. Leaves were placed side-by-side in a 12 cm^2^ (2×6 cm) leaf chamber (Li6400-11) attached to a LI-6400XT portable photosynthesis system (LiCor, Lincoln, NE, USA) fitted with a red-green-blue light (Li6400-18 RGB light source). At the time of measurement, all the plants were at tillering phase. During measurement, four different CO_2_ mole fractions in the leaf chamber (CO_2_ sample, ambient measurement CO_2_) were used (180, 280, 450 and 1000 μmol mol^−1^) with leaf temperature 25 °C, flow rate 350 μmol s^−1^ and irradiance at 1300 μmol m^−2^ s^−1^ using the red and blue LEDs only. Mesophyll conductance was measured using a stable carbon isotope tunable-diode laser absorption spectrometer (TDLAS, model TGA100A, Campbell Scientific, Inc., Logan, UT, USA) coupled to the gas exchange system as described by Barbour *et al*. (2007, 2010). The highest possible accuracy of gas exchange is required particularly for *A* and C_i_ from gas exchange to obtain reliable estimates of *g*_m_ (Pons *et al*., 2009). Similar CO_2_ concentrations around and inside the leaf chamber, and realistic values of *A* and C_i_ supported the conclusion that there were no leaks in this study that needed to be considered. Furthermore, Pons *et al*. (2009) observed that it is not necessary to check for leaks when net photosynthetic rates (*A*) are high (such as in wheat).

Mesophyll conductance was calculated from the difference between predicted discrimination assuming infinite mesophyll conductance (Δ_i_) and measured discrimination (Δ_obs_), as described by Jahan *et al*. (2014), using equations developed by Evans *et al*. (1986) and Barbour *et al*. (2010), and including a ternary effect as described by Farquhar and Cernusak (2012). Here, average daytime δ^13^C_growth_ δ^13^*C_CO2_* values were used for different ambient growth CO_2_ cabinets (−20.7, −18.4, −16.9 and −17.7 for 206, 344, 489 and 1085 μmol mol^−1^, respectively) in calculation of *g*_m_.

### Leaf Δ^13^C

Leaf samples of known area were collected directly after gas exchange measurement for Δ^13^C and percentage nitrogen concentration (%N). Leaves were oven dried at 65 °C for 72 hours and then dry mass was determined for LMA (leaf mass per area). Samples were then ground to a powder and approximately 1 mg of sample was weighed into tin cups. Isotope ratio mass spectrometry (IRMS) was used to determine the ratio of δ^13^C in samples. Samples were analysed on a DeltaZQ V Advantage coupled to a Conflo IV and a FlashHT in dual-reactor setup (Thermo Fisher Scientific, Bremen, Germany). The precision for the standard materials was between 0.04 ‰ and 0.09 ‰.

### Statistical analysis

The effects of CO_2_ concentration (mole fraction), drought treatment, and their interactions were assessed separately for both cultivars, and for each measure, namely *A, g*_sc_, *g*_m_ and *A*/*g*_sw_ (leaf-intrinsic water-use efficiency) using two-factor analysis of variance, as implemented in GenStat 14^th^ edition (VSN International Ltd, Hemel Hempstead, UK); means were compared using Fisher’s unprotected least significant difference tests. Differences were considered statistically significant when *P* < 0.05. Linear associations between the measures were evaluated using a general linear regression procedure in GenStat, making allowance for different regression slopes and intercepts between the two cultivars.

In order to explore all nine gas-exchange parameters simultaneously, a principal components analysis (PCA) was undertaken on these data (variables scaled to have a unit standard deviation), using the prcomp function in R (https://www.r-project.org/). Loadings of the first two principal components were interpreted, and this was also supported by calculation of Pearson correlation coefficients between all nine parameters.

## Results

### Growth CO_2_ effect

Plants were grown and measured at approximately the same CO_2_ mole fraction period (grown at 206, 344, 489 and 1085 μmol CO_2_ mol^−1^, and measured at 180, 280, 450 and 1000 μmol CO_2_ mol^−1^, respectively). Photosynthetic rate (*A*) was positively related (*P* < 0.001) to growth CO_2_ mole fraction when measured under saturating light and with the CO_2_ mole fraction surrounding the leaf controlled to match growth CO_2_ mole fraction, varying between 9 μmol m^−2^ s^−1^ (206 μmol CO_2_ mol^−1^) and 27 μmol m^−2^ s^−1^ (1085 μmol CO_2_ mol^−1^) (Fig. 1A & 1E). The two genotypes responded somewhat differently to growth CO_2_ mole fraction (i.e. there was a nearly-significant interaction effect between genotype and CO_2_; *P* = 0.06), with ‘Halberd’ grown at 489 μmol CO_2_ mol^−1^ having significantly higher *A* (23 μmol m^−2^ s^−1^) than ‘Cranbrook’ (18 μmol m^−2^ s^−1^) grown at the same CO_2_, despite the two genotypes having similar *A* at 1085 μmol CO_2_ mol^−1^. Irrigated plants had higher (*P* = 0.03) *A* compared to droughted plants.

**Fig. 1.**
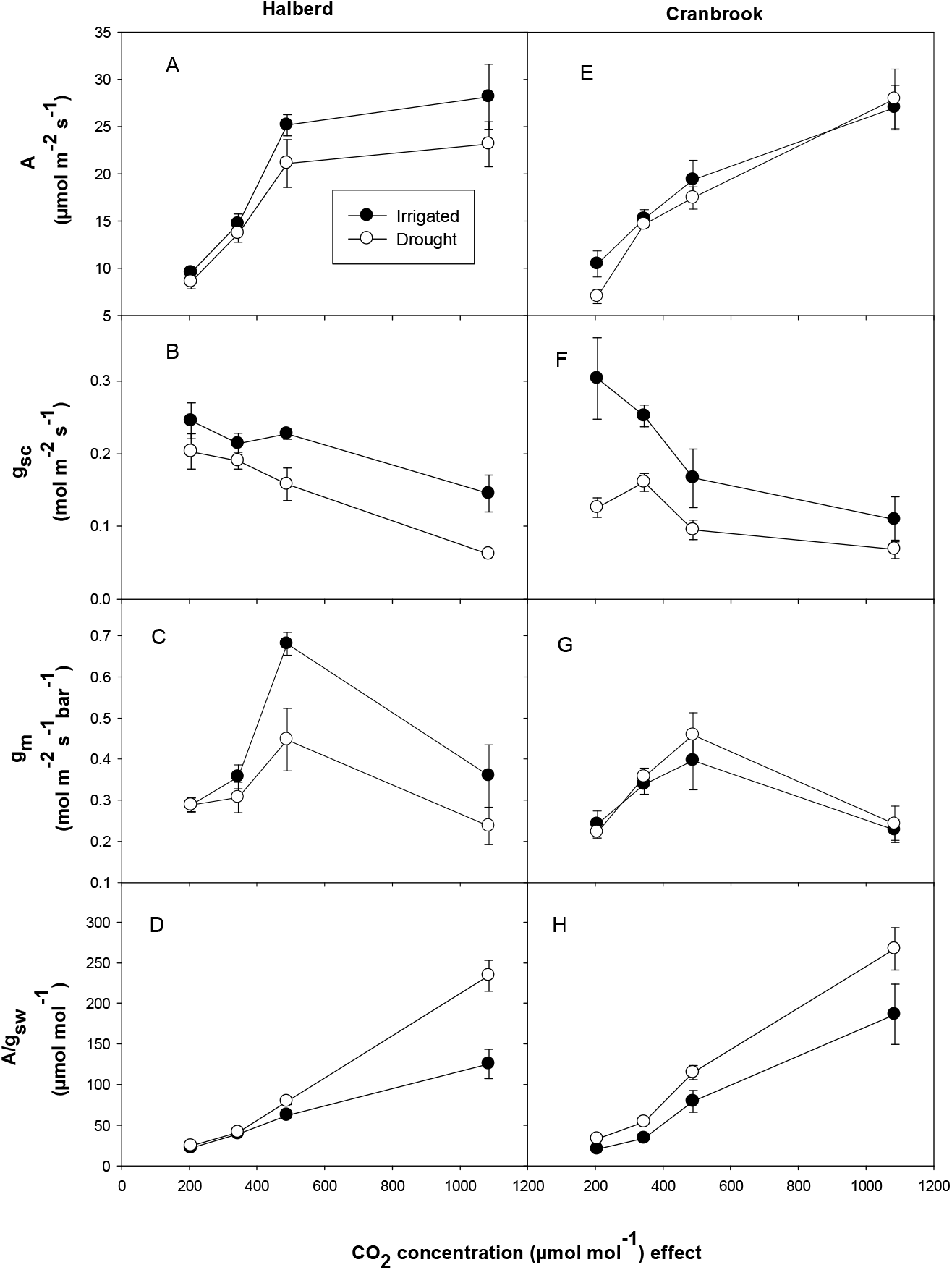
Growth [CO_2_] effect on photosynthetic rate (*A*), stomatal conductance to CO_2_ (*g*_sc_), mesophyll conductance (*g*_m_) and leaf-intrinsic water use efficiency (*A*/*g*_sw_) for leaves of two wheat cultivars. Values are mean ± standard error, *n* = 5.

Stomatal conductance (*g*_sc_) was negatively related to growth CO_2_ mole fraction (*P* < 0.001) when measured under saturating light and with the CO_2_ mole fraction surrounding the leaf controlled to match growth CO_2_ mole fraction, varying between 0.09 mol m^−2^ s^−1^ (1085 μmol CO_2_ mol^−1^) and 0.21 mol m^−2^ s^−1^ (206 μmol CO_2_ mol^−1^) (Fig. 1B & 1F). At any growth CO_2_ mole fraction, irrigated plants had significantly higher *g*_sc_ than droughted plants.

Mesophyll conductance (*g*_m_) was affected by growth CO_2_ mole fraction (*P* < 0.001), and increased with increasing CO_2_ concentration from 206 (0.26 mol m^−2^ s^−1^ bar^−1^) to 489 μmol CO_2_ mol^−1^ (0.50 mol m^−2^ s^−1^ bar^−1^), whereas at 1085 μmol CO_2_ mol^−1^, *g*_m_ was lower than 344 but higher than 206 μmol mol^−1^ (Fig. 1C & 1G). Irrigated plants had significantly higher *g*_m_ than droughted plants for ‘Halberd’ at 489 and 1085 growth CO_2_ mole fraction. Genotype ‘Halberd’ also had higher *g*_m_ than ‘Cranbrook’ at 489 μmol mol^−1^ growth CO_2_ mole fraction.

Leaf-intrinsic water-use efficiency (the ratio of photosynthetic rate to stomatal conductance of water vapour; *A*/*g*_sw_) increased significantly (*P* < 0.001) with increasing growth CO_2_ mole fraction from 206 to 1085 μmol mol^−1^ (Fig. 1C & 1G). At 1085 growth CO_2_ mole fraction, ‘Cranbrook’ had higher (*P* = 0.05) *A*/*g*_sw_ than ‘Halberd’. There was a significant interaction effect (*P* < 0.001) between CO_2_ and water availability on *A*/*g*_sw_, with increasingly higher *A*/*g*_sw_ for droughted plants compared to irrigated plants as CO_2_ mole fraction increased.

### Relationship of g_sc_ and g_m_ with A, and A/g_sw_ with g_m_/g_s_

Significant positive relationships (*P* < 0.001) were found between *g*_sc_ and *A*, and *g*_m_ and *A* at four different growth CO_2_ mole fractions (Fig. 2A & 2B). Between *g*_m_ and *A* (Fig. 2B), all the slopes were effectively equal, so that an increase in *g*_m_ is associated with the same increase in *A* at any of the four CO_2_ mole fraction. In pre-industrial and super-elevated [CO_2_] periods, the relationship between *g*_m_ and *A* was stronger (*r*^2^ = 0.66, and 0.62 respectively) than between *g*_sc_ and *A* (*r^2^* = 0.37, and 0.47 respectively). There was no significant water treatment effect on the association between *g*_sc_ and *A*, and between *g*_m_ and *A*. Leaf-intrinsic water-use efficiency (*A*/*g*_sw_) had a significant positive (*P* < 0.001) relationship with *g*_m_/*g*_sc_ for all four growth CO_2_ mole fractions periods (Fig. 3).

**Fig. 2.**
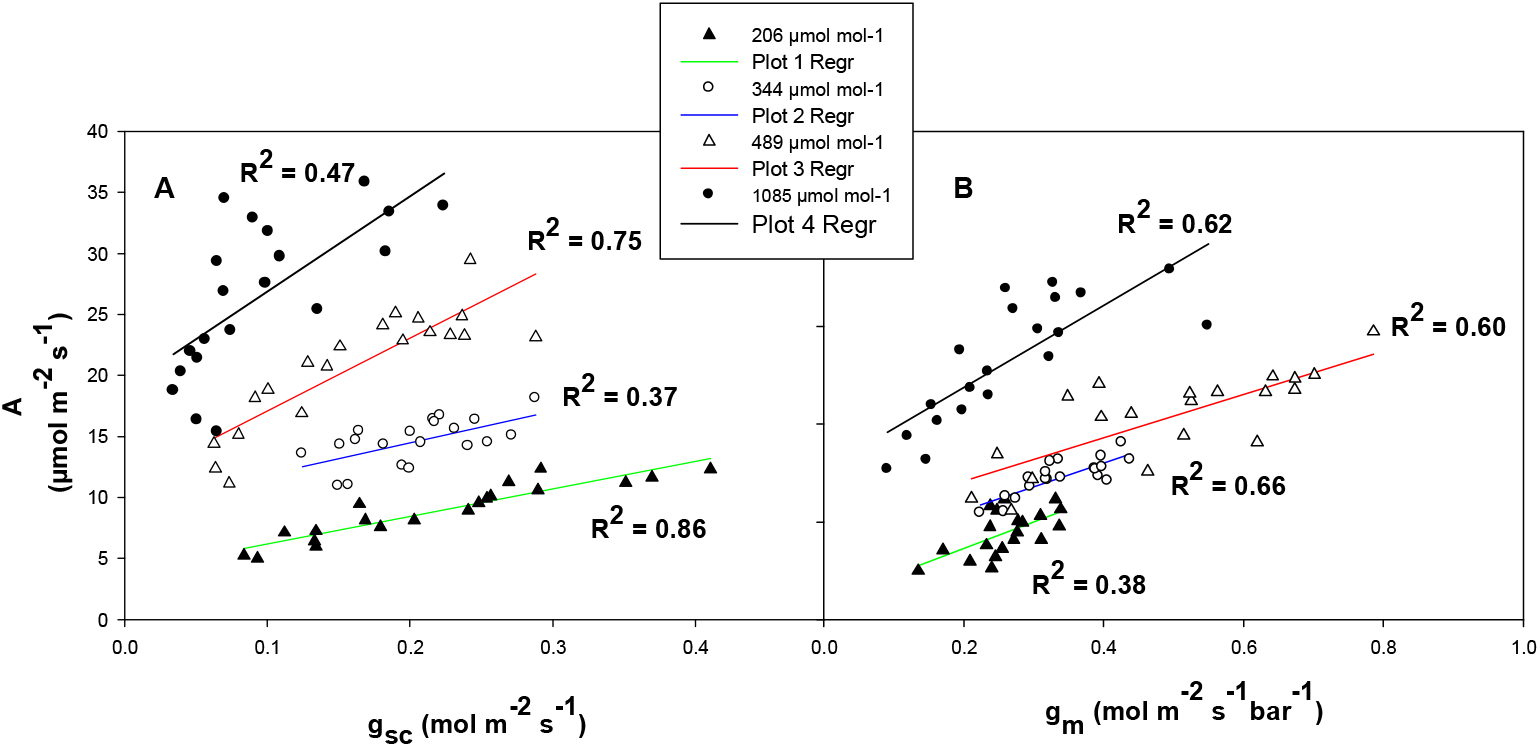
The relationships between photosynthetic rate and stomatal conductance to CO_2_ (A), and photosynthetic rate and mesophyll conductance (B) for both wheat genotypes at four different growth [CO_2_] (206, 344, 489 and 1085 μmol mol^−1^).

**Fig. 3.**
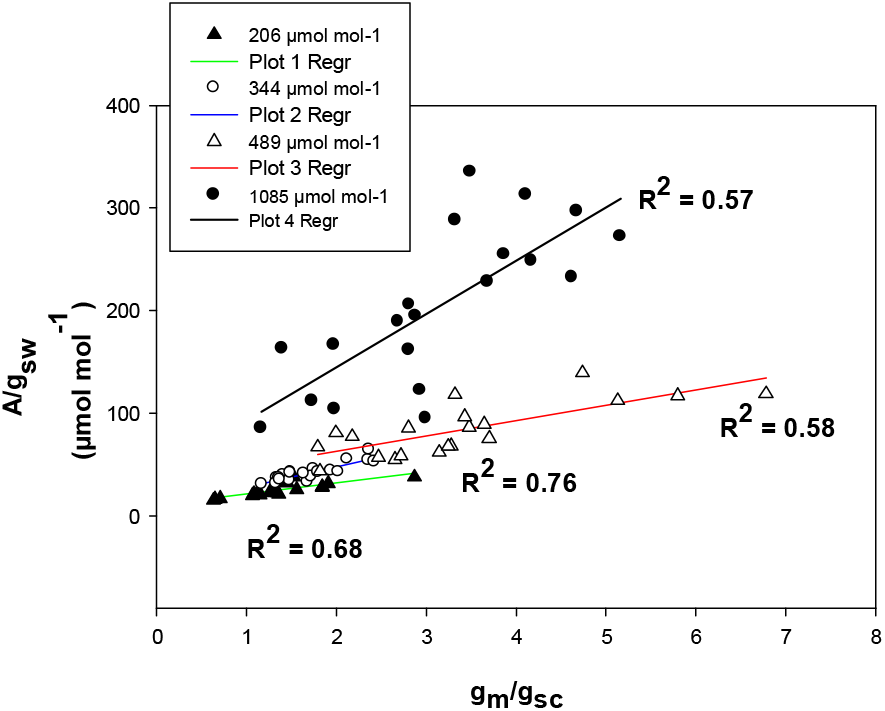
Correlation between leaf-intrinsic WUE (*A*/*g*_sw_) and the ratio of *g*_m_ and *g*_sc_ for both wheat genotypes at four different growth [CO_2_] (206, 344, 489 and 1085 μmol mol^−1^).

### Leaf Δ^13^C

The relationship between Δ^13^C and C_i_/C_a_ was not significant (*P* = 0.6, *r*^2^ = 0.024) (Fig. 4A), whereas the relationship between Δ^13^C and C_c_/C_a_ was significant (*P* = 0.04, *r*^2^ = 0.25) (Fig. 4B), when C_c_/C_a_ was calculated from gas exchange and online discrimination measurements made with the measurement CO_2_ mole fraction close to that of growth CO_2_ mole fraction. Moreover, there was a significant relationship between Δ C and g_m_ (*P* = 0.002, *r* = 0.5) (Fig. 5). Mesophyll conductance with %N had a positive relationship but not significant (*P* = 0.09, *r*^2^ = 0.18) (Fig. 6). Growth CO_2_ had a significant effect on LMA (*P* < 0.001) where LMA decreased from 204 to 489 and then increased at super elevated CO_2_ level (Fig. 7) however LMA did not show any significant relationships with *g*_m_ (Fig. 8).

**Fig. 4.**
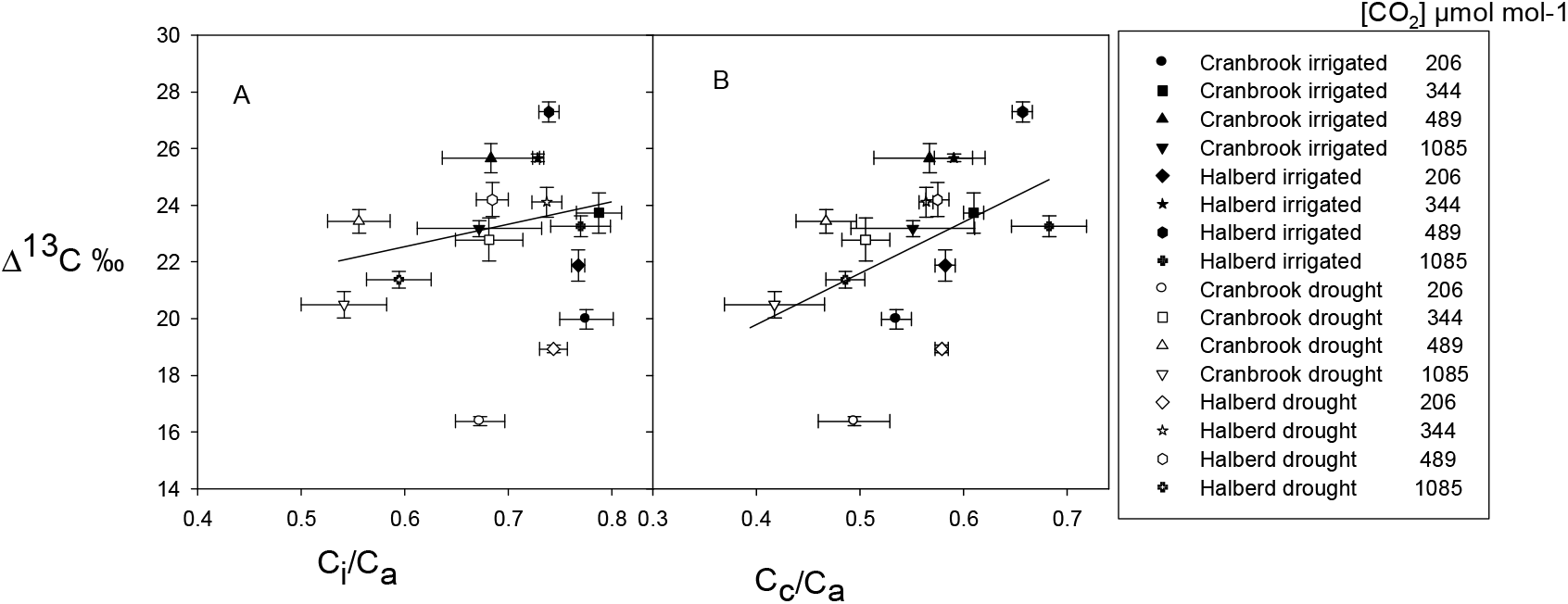
The relationship between Δ^13^C in leaf dry matter and the instantaneous ratios C_i_/C_a_ (A) and C_c_/C_a_ (B) in two wheat genotypes at four different growth CO_2_ mole fractions with two water treatments. Gas exchange parameters were measured at similar CO_2_ mole fraction as growth CO_2_. Values are means ± standard error, *n* = 5.

**Fig. 5.**
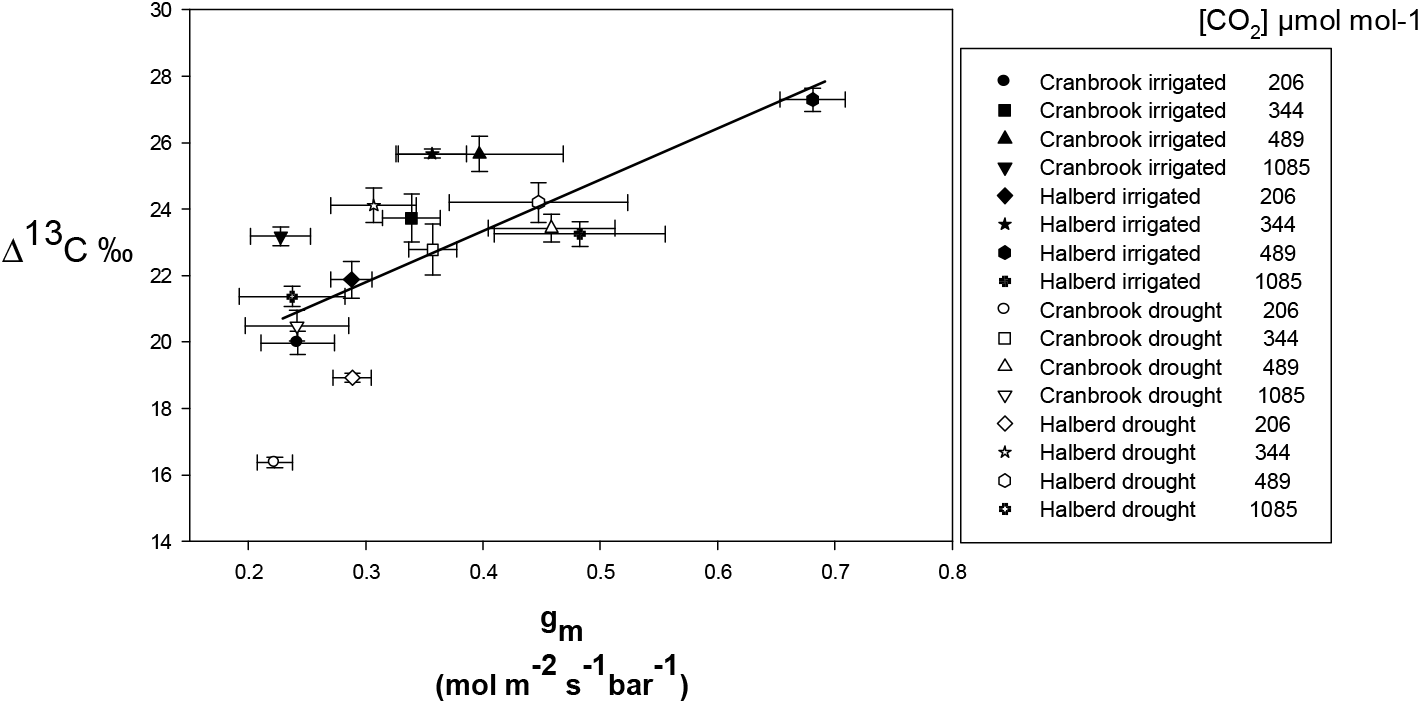
The relationships between mesophyll conductance and Δ^13^C in leaf dry matter for two wheat genotypes at four different growths CO_2_ mole fraction with two water treatments. The line is a least squares linear regression across all treatments: *g*_m_ = 0.031Δ^13^C - 0.37, *r*^2^ = 0.50, *P* = 0.002.

**Fig. 6.**
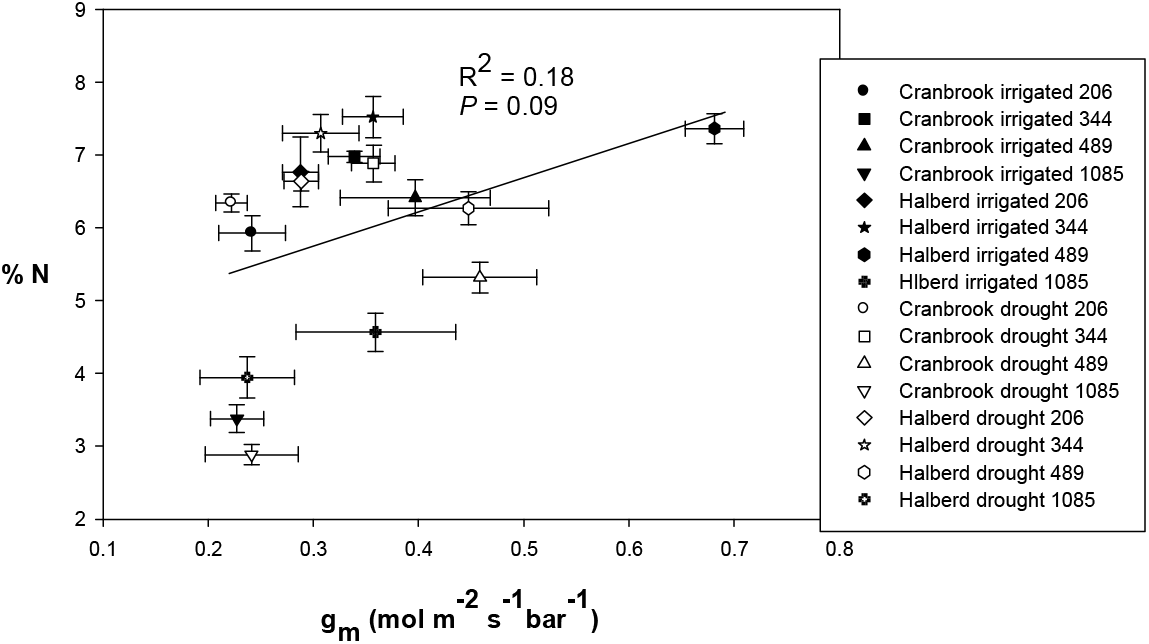
The relationship between % N in leaf dry matter and mesophyll conductance in two wheat genotypes at four different growth CO_2_ mole fractions with two water treatments. Gas exchange parameters were measured at similar CO_2_ mole fraction as growth CO_2_. Values are means ± standard error, *n* = 5, *r*^2^ = 0.15, *P* = 0.09.

**Fig. 7.**
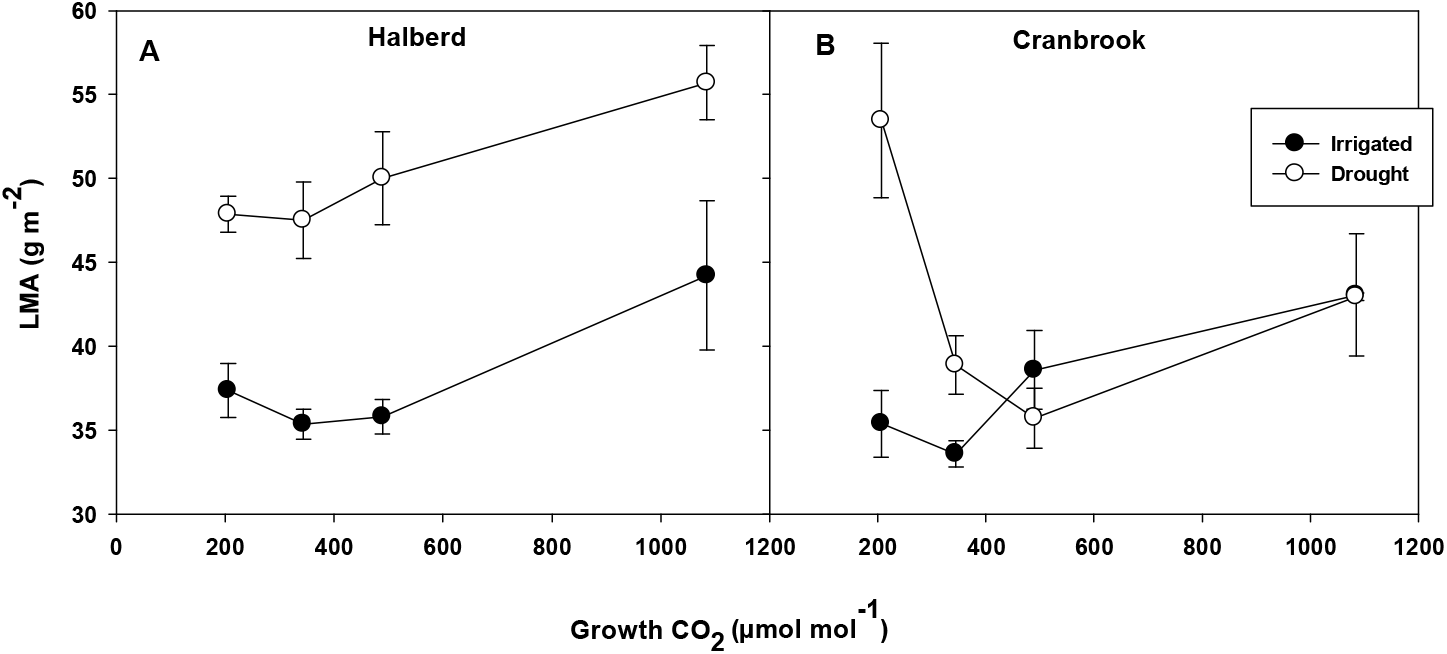
The response of leaf mass per area (LMA) at four growth CO_2_ mole fractions under two water treatments for genotype Halberd (A), and Cranbrook (B). Values are means ± standard error, *n* = 5.

**Fig. 8.**
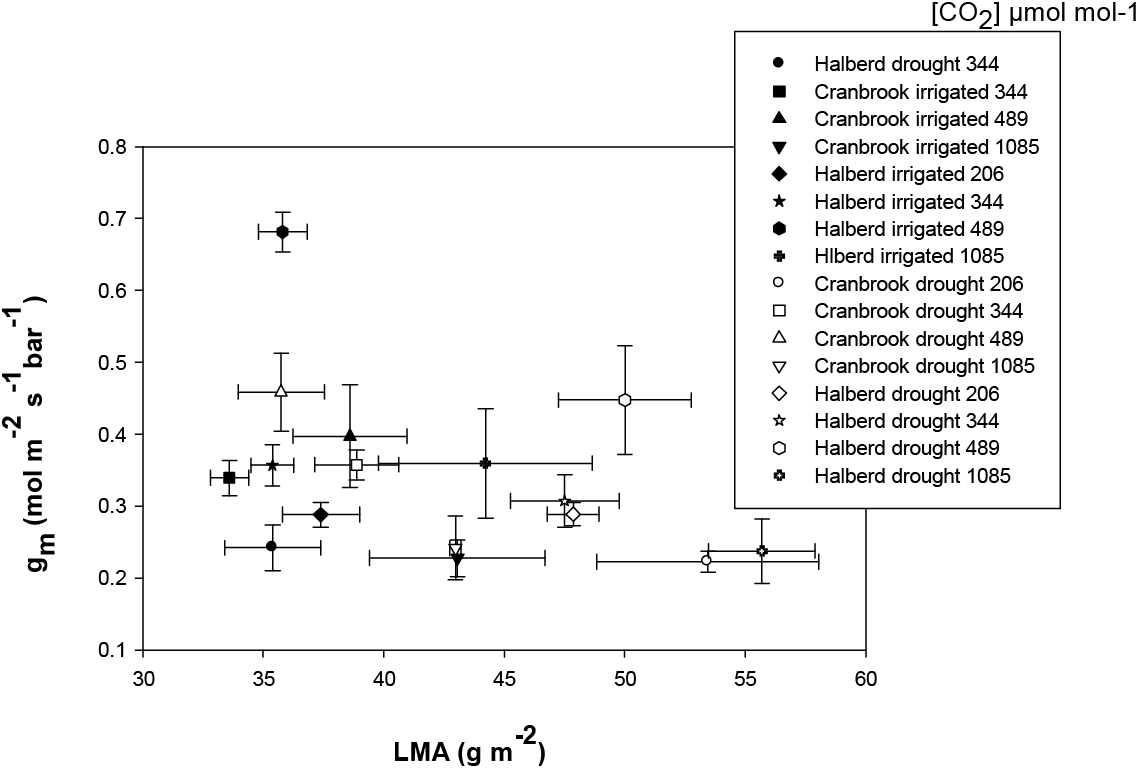
The relationship between leaf mass per area (LMA) and mesophyll conductance for two wheat genotypes at four different growths CO_2_ mole fraction with two water treatments. Values are means ± standard error, *n* = 5, *r*^2^ = 0.18, *P* = 0.13.

### Correlation between parameters and principal components analysis

The associations between all the parameters, including those mentioned above, can be summarised by correlations (Fig. 9). For example, the most positive correlation was between C_i_/C_a_ and C_c_/C_a_ (*r* = 0.84), while the most negative correlation was between %N and *A*/*g*_sw_ (*r* = −0.80). Exploring associations between all nine parameters further, based on the PCA, 68% of the variation in the nine-dimensional space was accounted for by the first two principal components (PC1 and PC2) (Table 1). In PC1, stomatal conductance (*g*_sc_), %N and the ratios C_c_/C_a_ and C_i_/C_a_ have high positive loadings, whereas *A*/*g*_sw_ has a high negative loading, so can be interpreted as a contrast between these first four parameters and *A*/*g*_sw_. On the other hand, in PC2, photosynthetic rate, mesophyll conductance and Δ^13^C have high negative loadings, and *A*/*g*_sw_ and C_c_/C_a_ have moderate negative loadings. From the plot of PC1 vs PC2, there is evidence of clustering of results on different growth [CO_2_] (Fig. 10).

**Fig. 9.**
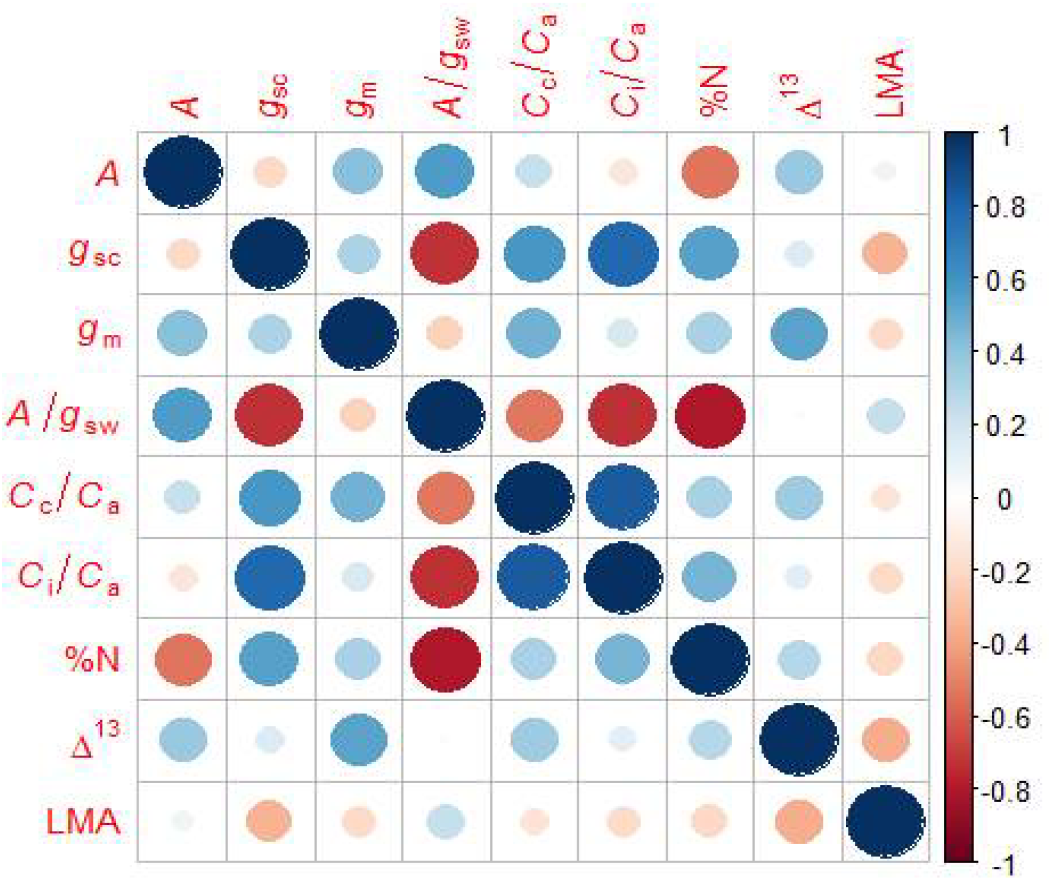
Visualization of the correlation matrix between all nine parameters.

**Table 1.** Principal component analysis and its component loadings.

|  | PC1 | PC2 |
| --- | --- | --- |
| Variation explained (%) | 44.2 | 23.4 |
| Variable |  |  |
| $A$ ($\mu\text{mol m}^{-2} \text{s}^{-1}$) | -0.12 | -0.61 |
| $g_{\text{sc}}$ ($\text{mol m}^{-2} \text{s}^{-1}$) | 0.43 | 0.05 |
| $g_{\text{m}}$ ($\text{mol m}^{-2} \text{s}^{-1} \text{bar}^{-1}$) | 0.23 | -0.45 |
| $A/g_{\text{sw}}$ ($\mu\text{mol mol}^{-1}$) | -0.44 | -0.27 |
| $C_{\text{c}}/C_{\text{a}}$ | 0.38 | -0.24 |
| $C_{\text{i}}/C_{\text{a}}$ | 0.42 | 0.04 |
| %N | 0.38 | 0.18 |
| $\Delta^{13}\text{C}$ | 0.17 | -0.47 |
| LMA | -0.17 | 0.11 |

**Fig. 10.**
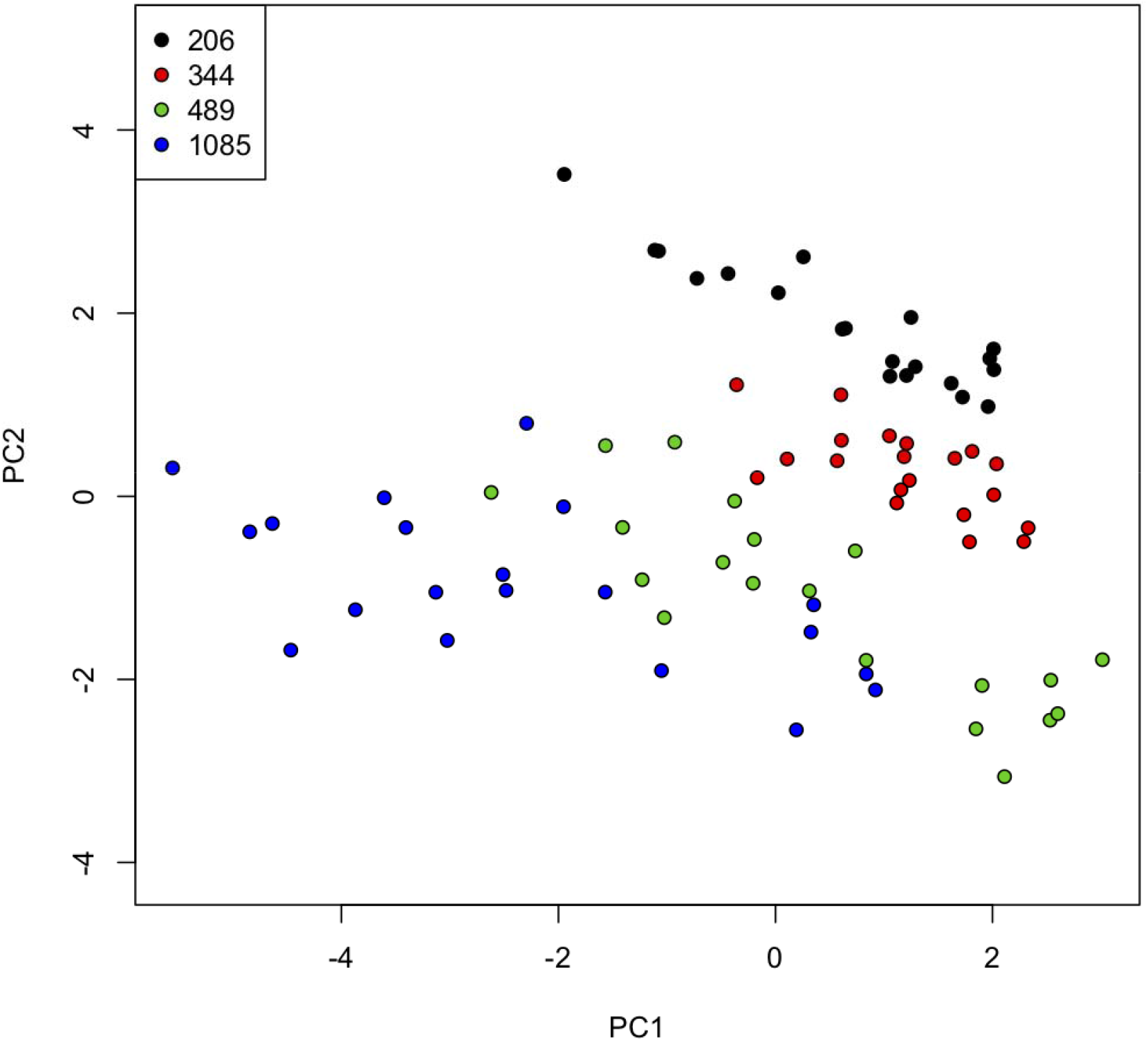
Plots of the first two principal component scores (PC1, PC2) for wheat plants. The growth [CO_2_] for each plant is indicated by the colour code.

## Discussion

### Growth [CO_2_]effects on g_m_

Mesophyll conductance increased with increasing growth [CO_2_] from 206 to 489 μmol mol^−1^ and then decreased at 1085 μmol mol^−1^ regardless of genotype and water treatment. Similarly, Singsaas *et al*. (2003) observed that *g*_m_ increased from 360 μmol mol^−1^ (ambient) to 560 μmol mol^−1^ CO_2_ concentration for some species [cucumber (*Cucumis sativus* L.), spinach (*Spinacia oleracea* L.) and linden bean (*Phaseolus vulgaris* L. var. *Linden*)] and then *g*_m_ decreased from 500 to 750 μmol mol^−1^ [CO_2_] for other species [sweetgum (*Liquidambar styraciflua* L.) and aspen (*Populus tremuloides* L.)]. Mesophyll conductance also decreased from 390 to 780 μmol mol^−1^ for *Arabidopsis thaliana* (Mizokami *et al*., 2019). In this study, the genotype ‘Halberd’ exhibitied a nearly two-fold higher *g*_m_ compared with ‘Cranbrook’ at ambient CO_2_ growth environment, which is similar to the results reported by Jahan *et al*. (2014). Moreover, ‘Halberd’ shows a higher response at super-elevated [CO_2_] compared to ‘Cranbrook’ (from Fig. 1C & 1G). Similarly, there was a species-dependent pattern on *g*_m_ with Ci ranging from 50 to 1500 μbar among six C3 species (depending on the species, *g*_m_ varied five-to nine-fold) (Flexas *et al*., 2007b). So, the response of *g*_m_ with different growth [CO_2_] also depends on species- and/or different environmental condition (Singsaas *et al*., 2003).

### The response of g_m_ has a physiological basis

During *g*_m_ calculation, we assumed values of two fractionation factors (*b* and *f*) and the non-photorespiratory respiration rate (*R*_d_), and these choices may possibly have altered *g*_m_ values with different [CO_2_]. For example, various estimates of fractionation associated with carboxylation (*b*) may be found in the literature between 26‰ and 31‰ (Brugnoli and Farquhar, 2000; McNevin *et al*., 2007; Lanigan *et al*., 2008) and the discrimination equation is particularly sensitive to *b* (Seibt *et al*., 2007). For this study, 29‰ for *b* was used for calculation. If a lower value of *b* is chosen (26‰; Lanigan *et al*., 2008), then □_i_ – □_obs_ becomes smaller and *g*_m_ larger, and if a higher value of *b* is chosen (31‰), then *g*_m_ becomes smaller. However, regardless of the value of *b* that is used, *g*_m_ is always higher for 489 μmol mol^−1^ and lower for 206 μmol mol^−1^.

*R_d_* values of 1.11 mol m^−2^ s^−1^ and 0.3 mol m^−2^ s^−1^ were used for genotypes ‘Cranbrook’ and ‘Halberd’, respectively for calculation of *g*_m_ (from Jahan *et al*., 2014). Jahan *et al*. (2014) performed a sensitivity analysis and showed that, at high light intensity, *g*_m_ estimates were not appreciably affected by the value used for *R*_d_. However, *R*_d_ was not measured at different CO_2_ mole fractions in this experiment, and sometimes *R*_d_ has been reported to be substantially decreased at high CO_2_ concentration (Bruhn *et al*., 2007). Gu and Sun (2014) mentioned that the use of an underestimate of the true *R*_d_ results in a strong nonlinear dependence of the estimated *g*_m_ on C_i_. If the *R*_d_ value was increased by 30% at a low CO_2_ mole fraction, then *g*_m_ will decrease slightly (from 0.34 to 0.32 mol m^−2^ s^−1^ bar^−1^ and from 0.40 to 0.39 mol m^−2^ s^−1^ bar^−1^ at 206 and 344 μmol mol^−1^, respectively. Similarly, if the *R*_d_ value is decreased by 30% at a high CO_2_ mole fraction (1000 μmol mol^−1^), then *g*_m_ will increase slightly (from 0.24 to 0.25 mol m s bar^−1^). So, estimates of *g*_m_ were not greatly affected by the value used for *R*_d_ at different CO_2_ mole fractions.

The value of 16.2‰ for fractionation during photorespiration (*f*) was used. Jahan *et al*. (2014) tested the sensitivity of calculated *g*_m_ to the values used for *f* by changing *f* from 16.2‰ to 11‰ for both 21% and 2% O_2_ concentrations, but did not find appreciable changes. However, it is possible that variable *f* could have an effect on the estimated values of *g*_m_ at different [CO_2_]. Cano *et al*. (2014) corrected the *g*_m_ estimation equation based on results reported by Tholen *et al*. (2012, 2014), which promoted the view that a low [CO_2_] increased the rate of photorespiration. If the highest *f* value (16.2‰) for 206 and 344 μmol mol^−1^ and the lowest value (11‰) for 1085 μmol mol^−1^ were used, the nonlinear relationship was also obtained with increasing *g*_m_ values from 206 to 1085 μmol mol^−1^. Flexas *et al*. (2007b) measured *g*_m_ at both 21% and 1% O2 with different Ci and found a decline of *g*_m_ at high CO_2_ for both 21% and 2% O2 levels; this suggests that variation during photorespiration can be neglected for *g*_m_ estimation. Moreover, if mitochondria are located predominately close to but behind the chloroplast, most of the photorespired CO_2_ may enter the chloroplast from the back, which makes the effect of photorespiration small or insignificant on the gradient between *p*_i_ and *p*_c_ (Cernusak *et al*., 2013). In one of our studies (unpublished), similar anatomical structures for wheat leaves were also observed when viewed under a light microscope, namely, that chloroplasts are located appressed to the cell wall with mitochondria located on the vascular side of the chloroplasts.

The reasons for *g*_m_ responses with [CO_2_] are varied. For example, Flexas *et al*. (2007b) worked on transgenic tobacco plants that differed in the amounts of aquaporin NtAQP1 and showed different slopes of *g*_m_ response to C_i_, and suggested that NtAQP1 may also be involved in these responses. Mizokami *et al*. (2019) suggested that abscisic acid (ABA) contents in leaves plays an important role in the regulation of *g*_m_ with [CO_2_] for *Arabidopsis thaliana*. Other studies have shown that *g*_m_ was limited by their mesophyll structure (Evans *et al*., 2009; Tosens *et al*., 2012b). In our study, LMA increased from glacial to ambient [CO_2_], and then decreased at super-elevated [CO_2_]; however, LMA was not correlated with *g*_m_ which may be due to the greater cell wall thickness which would constrain CO_2_ diffusion (Hassiotou *et al*., 2009; Tomas *et al*., 2013). There is another possibility that at super-elevated [CO_2_], some parts of the chloroplast are not adjacent to air spaces and this could lead to reduced *g*_m_ despite it having a high level of photosynthesis. Changes in chloroplast position and/or size could also affect *g*_m_ (Tholen *et al*., 2007). So, taken together there is evidence that the response of *g*_m_ with [CO_2_] has a physiological basis.

### Relationship of g_s_ and/or g_m_ with A, and improving WUE

The diffusion conductance of CO_2_ mostly includes *g*_s_ and *g*_m_, and it has been illustrated that *g*_s_ and *g*_m_ both were important determinants of *A* (Flexas *et al*., 2012; Tosen *et al*., 2012, 2016; Tomas *et al*., 2013). Additionally, this study observed that *g*_s_ and *g*_m_ are important determinants of *A* across all [CO_2_]. In the PCA, *g*_s_ and *g*_m_ also loaded highly on PC1 with concurring photosynthetic rates. So, both *g*_s_ and *g*_m_ play a critical role for achieving *A* not only for ambient [CO_2_] but also for future elevated [CO_2_], and neither *g*_s_ or *g*_m_ should be excluded in analyses of photosynthetic rate and carbon-cycle models. Despite highly variable values of *g*_m_, researchers have shown *g*_m_ to be the most limiting factor for *A* (Tomas *et al*., 2013; Tosen *et al*., 2016; Veromann-Jürgenson *et al*., 2017). Similarly, stronger associations of *g*_m_ with *A* compared to *g*_s_ with *A* were observed for simulated preindustrial and super-elevated [CO_2_]. Importantly, if *g*_s_ and *g*_m_ vary together, we need to understand how these traits affect resource use efficiency in a future changed climate.

In this study, from glacial to super-elevated [CO_2_], *g*_m_ increased with increasing *g*_sc_ (Fig. 11), and the correlation between *g*_m_ and *g*_sc_ also depended on [CO_2_]. There was an interesting pattern between *g*_m_ and *g*_sc_ at four different [CO_2_]; for a constant value of *g*_sc_, *g*_m_ increased with increasing CO_2_ mole fraction from 206 to 489 μmol mol^−1^ and then decreased at 1085 μmol mol^−1^ (Fig. 11); *g*_m_ was altered by different CO_2_ mole fractions to the same degree as *g*_sc_. The higher value of *g*_sc_ was not associated with a higher value of *g*_m_ at low growth CO_2_ mole fraction conditions. These relationships are supported by the PCA, where in PC1, *g*_sc_ had a high (positive) loading and *g*_m_ had a moderate (positive) loading.

**Fig. 11.**
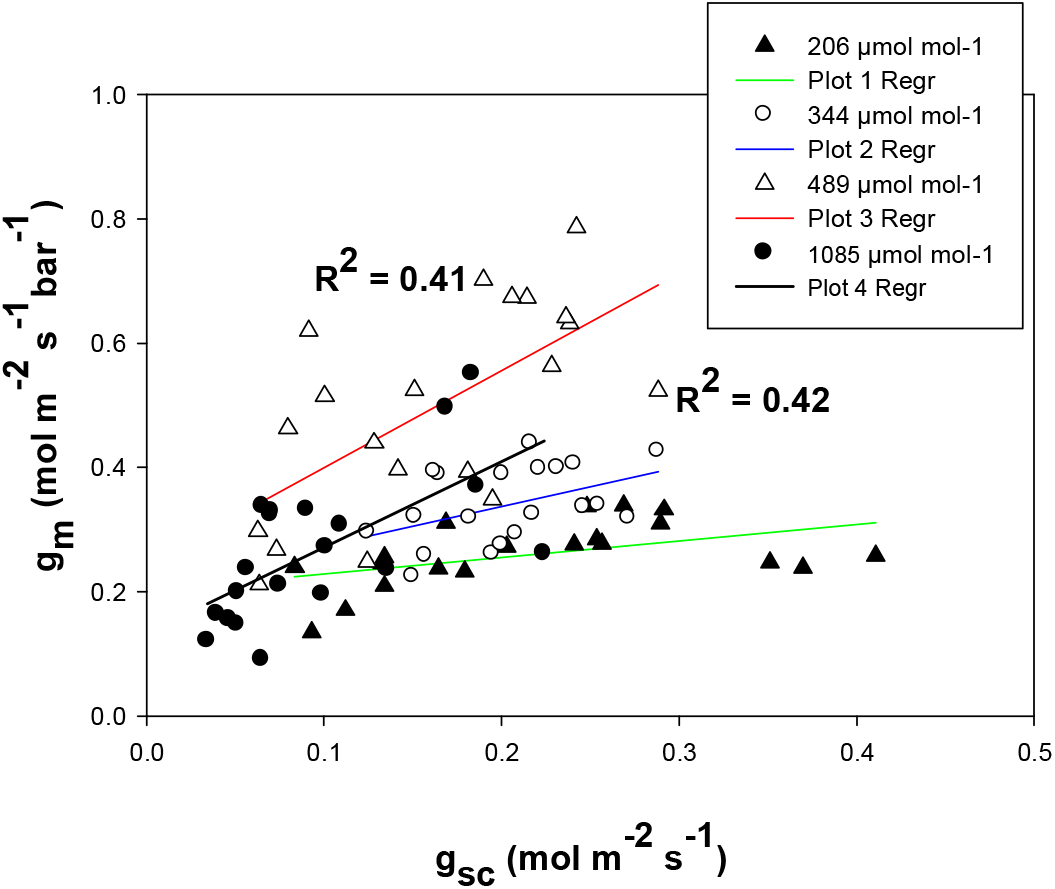
The relationships between stomatal conductance to CO_2_ and mesophyll conductance for both wheat genotypes at four different growth [CO_2_] (206, 344, 489 and 1085 μmol mol^−1^).

To improve WUE, it will be important to manage *g*_sc_, *g*_m_ and *g*_m_/*g*_sc_ simultaneously. Researchers observed that improving water-use efficiency through increased *g*_m_ would require *g*_sc_ and *g*_m_ to be uncoupled (Barbour *et al*., 2010; Gu *et al*., 2012; Jahan *et al*., 2014) because increasing *g*_m_ alone would result in increasing *A* which potentially increases leaf WUE. Moreover, researchers know that *g*_m_ can be affected by *g*_sc_, and the response of *g*_m_ to CO_2_ was faster than that of *g*_sc_ (Tazoe *et al*., 2011; Mizokami *et al*., 2019). In a water-stressed condition, *g*_m_ increased more than *g*_sc_ (Han *et al*., 2016), and this plays an important role in the change of *g*_m_/*g*_sc_. Our results show that leaf-intrinsic WUE (*A*/*g*_sw_) was strongly correlated with *g*_m_/*g*_sc_ for all four [CO_2_] periods (Fig. 3). We also observed that some of the increase in *g*_m_/*g*_sc_ was not associated with the same improvement rate of leaf-intrinsic WUE (*A*/*g*_sw_) for all growth [CO_2_] treatments; plants in the super-elevated treatment had higher rates of WUE than in the simulated glacial, pre-industrial or ambient treatments. Moreover, in this study we also have an interaction between the effect of growth CO_2_ and water treatment (here plants are only exposed to a mild water stress) on leaf-intrinsic WUE (*A*/*g*_sw_). Similarly, wheat plant water use efficiency (WUE) was 12% and 7% greater for Free-Air CO_2_ Enrichment (FACE) (200 μmol mol^−1^ above ambient) than the control environment (ambient 370 μmol mol^−1^ CO_2_) for first year and second year measurements, respectively (Hunsaker *et al*., 2000). Condon (2002) stated that field scale WUE (grain produced per unit water transpired) could be interpreted by leaf-intrinsic WUE. Barbour *et al*. (2010) stated that selecting for high *g*_m_ to improve *A*/*g*_sc_ will also improve crop WUE if allocation of carbon to the harvested plant organ is not reduced. Hence, for wheat genotypes, higher leaf-intrinsic WUE has the potential to improve crop WUE.

The carbon isotope composition of leaf tissue strongly reflects photosynthetic discrimination (Farquhar *et al*., 1989), so the ratio of chloroplast to ambient CO_2_ mole fractions, C_c_/C_a_, should be taken into account for evaluating *A, g*_sc_ and *g*_m_. The observation here of no significant correlation between C_i_/C_a_ and Δ^13^C, but a significant correlation between C_c_/C_a_ and Δ^13^C supports this theory. Δ^13^C in leaf dry matter has been widely used as a long-term estimation method for C_i_/C_a_ and WUE between species or genotypes (Brugnoli and Farquhar, 2000), assuming either constant *g*_m_, or that *g*_m_ scales with *g*_sc_. Moreover, ^13^C discrimination (Δ) and %N in leaf dry matter also shows a similar trend to that observed in *g*_m_ (increased with increasing growth CO_2_ from 206 to 489, then decreased at 1085 μmol mol^−1^). Therefore, *g*_m_ plays an important role in regulating *A*, which could alter WUE for stressed crops in a future environment.

## Conclusion

In two wheat genotypes, *g*_m_ increased with increasing [CO_2_] from simulated glacial era to ambient condition and then declined at super-elevated [CO_2_]. Although the specific mechanism of *g*_m_ response to [CO_2_] remains unclear, the response of *g*_m_ to [CO_2_] may be due to aquaporins (Flexas *et al*., 2007b) and/or leaf anatomical changes (Tholen *et al*., 2007) which have a physiological basis. However, *g*_m_ and *g*_m_/*g*_sc_ are strongly associated with the variability of *A* and leaf-intrinsic WUE, especially for future super-elevated [CO_2_]. Moreover, *g*_m_ may be more important than *g*_sc_ in the change of *g*_m_/*g*_sc_, so plants with higher *g*_m_ may improve *A* without increasing *g*_sc_, which ultimately increases leaf-intrinsic WUE (*A*/*g*_sw_) for a climactically changed future environment. These data will inform plant breeding programs for Australian wheat under future environmental conditions. Additional gains could also be made by searching for genomic regions associated with these WUE-related traits through quantitative trait loci (QTL) or genome-wide association study (GWAS). Such a breeding program may then be further enhanced by inclusion of this genomic information in their selection schemes.

## Acknowledgements

Firstly, I would like to express my sincere thanks to my PhD supervisor, Professor Margaret Barbour, who gave support throughout my experiment and editorial assistance. I thank Svetlana Ryazanova for technical assistance with gas exchange and stable isotope measurements. This work was supported by the Grains Research and Development Corporation (US00056, and GRS 10660 to support my study).

